# Genomic epidemiology of the current wave of artemisinin resistant malaria

**DOI:** 10.1101/019737

**Authors:** Roberto Amato, Olivo Miotto, Charles Woodrow, Jacob Almagro-Garcia, Ipsita Sinha, Susana Campino, Daniel Mead, Eleanor Drury, Mihir Kekre, Mandy Sanders, Alfred Amambua-Ngwa, Chanaki Amaratunga, Lucas Amenga-Etego, Tim J C Anderson, Voahangy Andrianaranjaka, Tobias Apinjoh, Elizabeth Ashley, Sarah Auburn, Gordon A Awandare, Vito Baraka, Alyssa Barry, Maciej F Boni, Steffen Borrmann, Teun Bousema, Oralee Branch, Peter C Bull, Kesinee Chotivanich, David J Conway, Alister Craig, Nicholas P Day, Abdoulaye Djimdé, Christiane Dolecek, Arjen M Dondorp, Chris Drakeley, Patrick Duffy, Diego F Echeverri-Garcia, Thomas G Egwang, Rick M Fairhurst, Abul Faiz, Caterina I Fanello, Tran Tinh Hien, Abraham Hodgson, Mallika Imwong, Deus Ishengoma, Pharath Lim, Chanthap Lon, Jutta Marfurt, Kevin Marsh, Mayfong Mayxay, Victor Mobegi, Olugbenga Mokuolu, Jacqui Montgomery, Ivo Mueller, Myat Phone Kyaw, Paul N Newton, Francois Nosten, Rintis Noviyanti, Alexis Nzila, Harold Ocholla, Abraham Oduro, Marie Onyamboko, Jean-Bosco Ouedraogo, Aung Pyae Phyo, Christopher V Plowe, Ric N Price, Sasithon Pukrittayakamee, Milijaona Randrianarivelojosia, Pascal Ringwald, Lastenia Ruiz, David Saunders, Alex Shayo, Peter Siba, Shannon Takala-Harrison, Thuy-Nhien Nguyen Thanh, Vandana Thathy, Federica Verra, Nicholas J White, Ye Htut, Victoria J Cornelius, Rachel Giacomantonio, Dawn Muddyman, Christa Henrichs, Cinzia Malangone, Dushyanth Jyothi, Richard D Pearson, Julian C Rayner, Gilean McVean, Kirk Rockett, Alistair Miles, Paul Vauterin, Ben Jeffery, Magnus Manske, Jim Stalker, Bronwyn MacInnis, Dominic P Kwiatkowski, MalariaGEN *Plasmodium falciparum* Community Project

## Abstract

Artemisinin resistant *Plasmodium falciparum* is advancing across Southeast Asia in a soft selective sweep involving at least 20 independent *kelch13* mutations. In a large global survey, we find that *kelch13* mutations which cause resistance in Southeast Asia are present at low frequency in Africa. We show that African *kelch13* mutations have originated locally, and that *kelch13* shows a normal variation pattern relative to other genes in Africa, whereas in Southeast Asia there is a great excess of non-synonymous mutations, many of which cause radical amino-acid changes. Thus, *kelch13* is not currently undergoing strong selection in Africa, despite a deep reservoir of standing variation that could potentially allow resistance to emerge rapidly. The practical implications are that public health surveillance for artemisinin resistance should not rely on *kelch13* data alone, and interventions to prevent resistance must account for local evolutionary conditions, shown by genomic epidemiology to differ greatly between geographical regions.

## Introduction

Artemisinin combination therapy (ACT), the frontline treatment for *P. falciparum* infection, has played a major part in reducing the number of deaths due to malaria over the past decade (World Health Organization 2014). However artemisinin resistant *P. falciparum*, which has recently spread across large parts of Southeast Asia, now threatens to destabilise malaria control worldwide (Dondorp et al. 2009; Hien et al. 2012; Phyo et al. 2012; Kyaw et al. 2013; Ashley et al. 2014; World Health Organization 2014). One of the main contemporary challenges in global health is to prevent artemisinin resistance from becoming established in Africa, where the consequences for childhood mortality could be disastrous (Dondorp and Ringwald 2013).

Understanding the epidemiological and evolutionary processes that are driving the current wave of artemisinin resistance is essential to develop effective strategies to stop it spreading. At the molecular level, artemisinin resistance is caused by mutations in a kelch protein encoded by PF3D7_1343700 on *P. falciparum* chromosome 13, referred to here as *kelch13*. More specifically, non-synonymous mutations in the *kelch13* propeller and BTB-POZ domains (KPBD) result in reduced sensitivity of *P. falciparum* to artemisinin, as demonstrated by multiple lines of evidence including laboratory studies of artificially acquired resistance, genetic association studies of natural resistance and allelic replacement experiments (Ariey et al. 2014; Ghorbal et al. 2014; Miotto et al. 2015; Straimer et al. 2015; Takala-Harrison et al. 2015). The precise biological function of *kelch13* is still not fully understood, but parasites with KBPD mutations tend to grow more slowly in the early part of the erythrocytic cycle, and have an enhanced unfolded protein response, both of which might act to protect against oxidative damage caused by artemisinin (Dogovski et al. 2015; Mbengue et al. 2015; Mok et al. 2015).

A striking characteristic of the current wave of artemisinin resistance is that it is caused by multiple independent KPBD mutations emerging in different locations, i.e. it does not originate from a single mutational event. More than 20 KPBD SNPs have been associated with delayed parasite clearance during artemisinin treatment and there are several documented instances of the same allele arising independently in different locations (Ashley et al. 2014; Miotto et al. 2015; Takala-Harrison et al. 2015). These are classic features of a soft selective sweep which, according to evolutionary theory, is most likely to arise in large populations where the selected alleles are already present as standing genetic variation (Hermisson and Pennings 2005; Pennings and Hermisson 2006; Messer and Petrov 2013). There is ongoing debate among evolutionary biologists about how commonly soft selective sweeps occur in nature (Jensen 2014; Garud et al. 2015) but they have clearly played a role in previous forms of antimalarial drug resistance (Nair et al. 2007; Salgueiro et al. 2010) and the current wave of artemisinin resistance is the most extreme example of a soft selective sweep thus far observed in eukaryotes.

This creates a practical problem in monitoring the global spread of resistance. Artemisinin resistance can be measured directly, by following the rate of parasite clearance in patients (Flegg et al. 2011) or by testing parasite isolates *in vitro* (Witkowski et al. 2013), but these phenotypic assays are resource intensive and impractical for large-scale screening in resource-poor settings. Genetic approaches are therefore preferable for practical implementation of large-scale surveillance, but the soft selective sweep of artemisinin resistance produces much more heterogeneous genetic signatures than previous global waves of chloroquine and pyrimethamine resistance, where hard selective sweeps were the dominant mode of spread. Thus there is considerable uncertainty about the epidemiological significance of the growing number of non-synonymous KPBD mutations reported in Africa (Ashley et al. 2014; Hopkins Sibley 2015; Kamau et al. 2015; Taylor et al. 2015). Previous studies in Africa have not identified variants known to cause resistance in Southeast Asia (Kamau et al. 2015; Taylor et al. 2015), and there are documented instances of African parasites with KPBD mutations clearing rapidly following artemisinin treatment (Ashley et al. 2014), but in the absence of comprehensive phenotypic data it is not known which if any of these mutations are markers of resistance. This is a limitation of conventional molecular epidemiology, which tracks specific mutations and haplotypes and is poorly equipped to monitor soft selective sweeps where new mutations are continually arising on different haplotypic backgrounds, making it difficult to keep track of their phenotypic effects and evolutionary trajectories.

Here we explore how genomic epidemiology might help overcome these practical obstacles to monitoring the current wave of artemisinin resistance. This analysis includes genome sequencing data for 3,411 clinical samples of *P. falciparum* obtained from 46 locations in 23 countries. This large dataset was generated by the MalariaGEN *Plasmodium falciparum* Community Project, a collaborative study in which multiple research groups working on different scientific questions are sharing genome variation data to generate an integrated view of polymorphism in the global parasite population.

## Results

### Africa and Southeast Asia both have many *kelch13* polymorphisms

Paired-end sequence reads were generated using the Illumina platform and aligned to the *P. falciparum* 3D7 reference genome, applying a series of quality control filters as previously described (see Methods) (Manske et al. 2012; Miotto et al. 2013). The initial alignments identified 4,305,957 potential SNPs, which after quality control filtering produced a set of 926,988 exonic SNPs that could be genotyped with high confidence in the majority of samples, and that are used as the basis for this analysis.

As summarised in Table 1, the dataset comprised 1,648 samples from Africa and 1,599 samples from Southeast Asia, allowing us to compare these two groups directly without the need for sample size corrections. We identified a total of 155 SNPs in the *kelch13* gene, of which 128 were seen in Africa and 62 in Southeast Asia (Table 2). Studies in Southeast Asia have found that artemisinin resistance is associated with non-synonymous polymorphisms in the propeller and adjacent BTB/POZ domains, collectively referred to here as the *kelch13* resistance domain. Out of a total of 46 non-synonymous SNPs in the resistance domain (Table 3) we found a similar number in Africa (n=26) and Southeast Asia (n=34), with 14 seen in both places. Seven of those observed in Africa have previously been associated with artemisinin resistance in Southeast Asia, including C580Y, the most common allele in resistant parasites (Miotto et al. 2015).

**Table 1.** Count of samples analysed in this study.

| Region | Code | Sample counts | Country | Code | Sample counts |
| --- | --- | --- | --- | --- | --- |
| **West Africa** | WAF | 957 | Burkina Faso | BF | 56 |
|  |  |  | Cameroon | CM | 134 |
|  |  |  | Ghana | GH | 478 |
|  |  |  | Gambia, The | GM | 73 |
|  |  |  | Guinea | GN | 124 |
|  |  |  | Mali | ML | 87 |
|  |  |  | Nigeria | NG | 5 |
| **East Africa** | EAF | 412 | Kenya | KE | 52 |
|  |  |  | Madagascar | MG | 18 |
|  |  |  | Malawi | MW | 262 |
|  |  |  | Tanzania | TZ | 68 |
|  |  |  | Uganda | UG | 12 |
| **Central Africa** | CAF | 279 | D. R. Congo | CD | 279 |
| **South America** | SAM | 27 | Colombia | CO | 16 |
|  |  |  | Peru | PE | 11 |
| **South Asia** | SAS | 75 | Bangladesh | BD | 75 |
| **West Southeast Asia** | WSEA | 497 | Myanmar | MM | 111 |
|  |  |  | Thailand | TH | 386 |
| **East Southeast Asia** | ESEA | 1102 | Cambodia | KH | 762 |
|  |  |  | Laos | LA | 120 |
|  |  |  | Vietnam | VN | 220 |
| **Oceania** | OCE | 62 | Indonesia (Papua) | ID | 17 |
|  |  |  | Papua New Guinea | PG | 45 |

**Table 2.** Worldwide distribution of *kelch13* mutations. Number of mutations present (not necessarily exclusively) in 5 populations (AFR=Africa, SEA=Southeast Asia, SAS=South Asia, OCE=Oceania, SAM=South America).

|  |  | AFR | SEA | SAS | OCE | SAM |
| --- | --- | --- | --- | --- | --- | --- |
| *Non-synonymous* | *Resistance domains* | 26 | 34 | 1 | 1 | 0 |
|  | *Upstream region* | 42 | 16 | 2 | 1 | 1 |
| *Synonymous* | *Resistance domains* | 38 | 9 | 1 | 0 | 0 |
|  | *Upstream region* | 22 | 3 | 1 | 0 | 0 |

**Table 3.** List of the non-synonymous mutations in the *kelch13* resistance domains. Non-synonymous mutations found in the *kelch13* resistance domains and their position on chromosome Pf3D7_13_v3. For each mutation is reported where it has been observed and in how many samples. Where known, we reported if the mutation has been previously observed in patients with a prolonged parasite clearance half-life (>5 hours) by Miotto et al. 2014 and/or Ashley et al 2014.

| Mutation | Genomic coordinates | AFR | SEA | SAS | OCE | SAM | Observed in ART-R samples? |
| --- | --- | --- | --- | --- | --- | --- | --- |
| **D353Y** | 1725941 | - | **4** | - | - | - | **Yes** |
| **F395Y** | 1725814 | - | 1 | - | - | - | No |
| **I416V** | 1725752 | 1 | 1 | - | - | - |  |
| **I416M** | 1725750 | 1 | - | - | - | - |  |
| **K438N** | 1725684 | - | 1 | - | - | - | No |
| **P441L** | 1725676 | - | **27** | - | - | - | **Yes** |
| **P443S** | 1725671 | - | 1 | - | - | - |  |
| **F446I** | 1725662 | - | **7** | - | - | - | **Yes** |
| **G449A** | 1725652 | - | **7** | - | - | - | **Yes** |
| **S459L** | 1725622 | 2 | 2 | - | - | - |  |
| **A481V** | 1725556 | - | **4** | - | - | - | **Yes** |
| **S485N** | 1725544 | - | 1 | - | - | - |  |
| **Y493H** | 1725521 | **1** | **76** | - | - | - | **Yes** |
| **V520I** | 1725440 | 1 | - | - | - | - |  |
| **S522C** | 1725434 | 2 | 1 | - | - | - | No |
| **P527H** | 1725418 | 1 | 5 | - | - | - |  |
| **C532S** | 1725404 | 1 | - | - | - | - |  |
| **V534L** | 1725398 | 2 | - | - | - | - |  |
| **N537I** | 1725388 | 1 | 1 | - | - | - | No |
| **G538V** | 1725385 | - | **19** | - | - | - | **Yes** |
| **R539T** | 1725382 | - | **63** | - | - | - | **Yes** |
| **I543T** | 1725370 | - | **34** | - | - | - | **Yes** |
| **P553L** | 1725340 | **2** | **24** | - | - | - | **Yes** |
| **A557S** | 1725329 | 1 | - | - | - | - |  |
| **R561H** | 1725316 | **1** | **24** | - | - | - | **Yes** |
| **V568G** | 1725295 | - | **6** | - | - | - | **Yes** |
| **T573S** | 1725280 | 2 | - | - | - | - |  |
| **P574L** | 1725277 | - | **12** | - | - | - | **Yes** |
| **R575K** | 1725274 | - | 3 | - | - | - |  |
| **A578S** | 1725266 | 18 | - | - | - | - | No |
| **C580Y** | 1725259 | **2** | **423** | - | **1** | - | **Yes** |
| **D584V** | 1725247 | - | **3** | - | - | - | **Yes** |
| **V589I** | 1725233 | 2 | - | - | - | - |  |
| **T593S** | 1725221 | 1 | - | - | - | - |  |
| **E612D** | 1725162 | 1 | - | - | - | - |  |
| **Q613E** | 1725161 | 5 | 1 | - | - | - |  |
| **Q613L** | 1725160 | 1 | - | - | - | - | No |
| **F614L** | 1725158 | - | 1 | - | - | - | No |
| **Y630F** | 1725109 | 2 | 1 | - | - | - |  |
| **V637I** | 1725089 | 2 | - | - | - | - |  |
| **P667A** | 1724999 | - | 2 | - | - | - |  |
| **P667L** | 1724998 | - | 2 | 1 | - | - |  |
| **F673I** | 1724981 | - | **3** | - | - | - | **Yes** |
| **A675V** | 1724974 | **1** | **18** | - | - | - | **Yes** |
| **A676S** | 1724972 | 2 | 3 | - | - | - |  |
| **H719N** | 1724843 | **1** | **8** | - | - | - | **Yes** |

### *kelch13* polymorphisms in Africa appear to be indigenous

We asked whether kelch13 polymorphisms seen in Africa had emerged independently, or they had migrated from Southeast Asia. We started by grouping samples according to genome-wide genetic similarity, based on a neighbour-joining (NJ) tree (Figure 1). African and Southeast Asian samples formed two well-separated and distinct clusters, suggesting that gene flow between the two regions is very modest or negligible. None of the African parasites carrying *kelch13* mutations grouped with the Southeast Asian population, supporting the idea that these mutations are indigenous.

**Figure 1.**
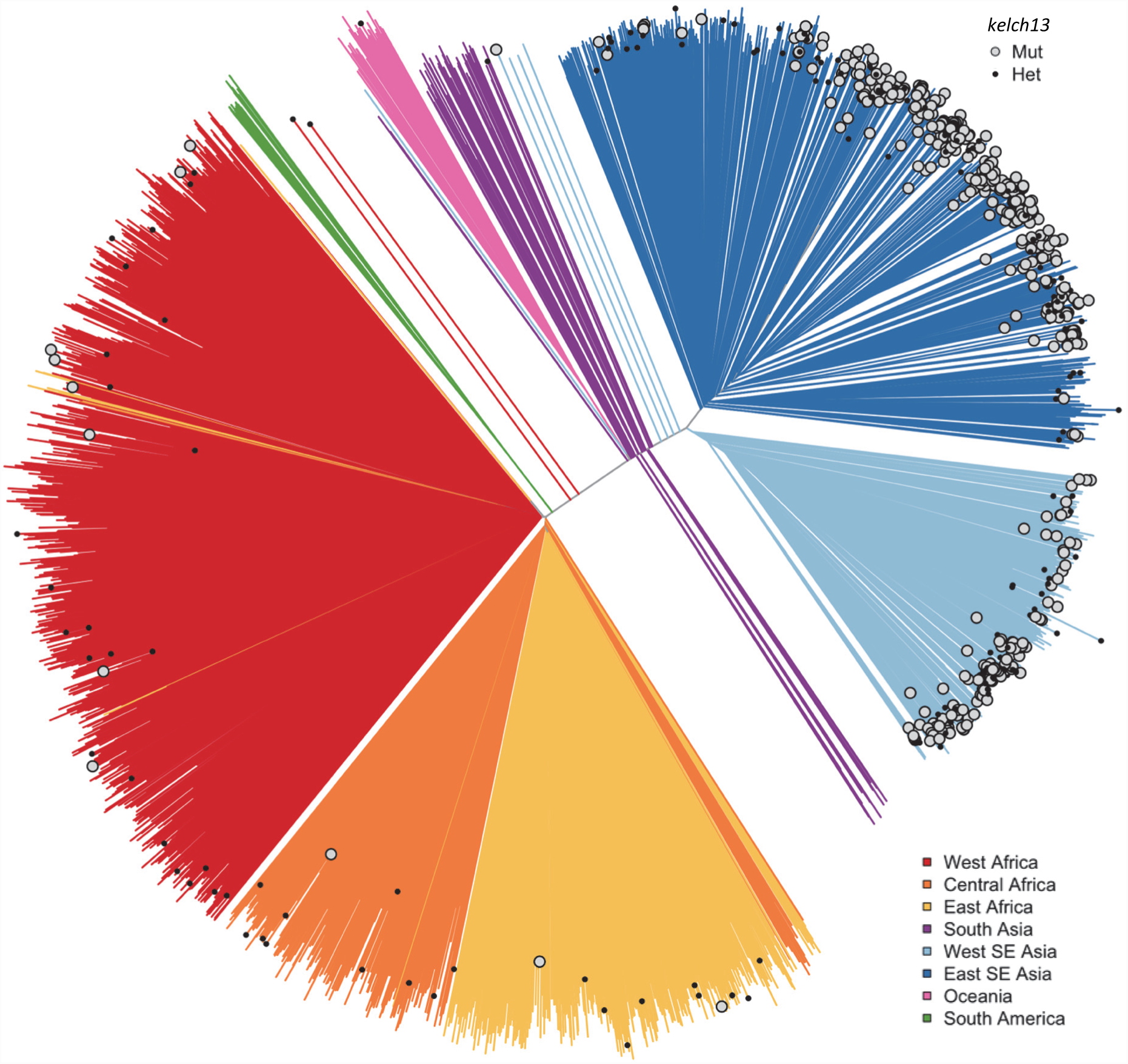
Different origin of African and Asian mutations. Neighbour-joining tree showing population structure across all the sampling sites. Countries are classified as in Table 1. Branches with large open tip circle indicate the sample is a *kelch13* mutant, while those with a small black symbol are mixed infections (i.e. mixture of wild-type and mutant parasites or two mutant parasites with different mutations). Branches without tip symbols are *kelch13* wild type. African *kelch13* mutants are, at a genomic level, more similar to other African samples.

Two African *kelch13* mutants did not cluster with the bulk of African samples, but occupied a somewhat isolated position in the NJ tree. We note that these two samples exhibit unusually high levels of heterozygosity (F_WS_ < 0.4), with mixed calls randomly distributed across the genome, and no evidence of recent recombination events. Continuous genetic monitoring of the parasite population will determine whether these are indeed just isolated cases, or they constitute very early evidence of gene flow between the two regions; at this stage we cannot rule out that they are more simply the result of biological contamination during preparation and processing.

Genome-wide analysis cannot identify gene flow from Southeast Asia which occurred in earlier times and, after recombination with local haplotypes, resulted in the transfer of DNA segments into genomes that otherwise appear to be local. Thus, we analyzed regions flanking *kelch13* in a 100kbp genomic region around the gene, to detect fragments of non-African haplotypes that might have remained in linkage disequilibrium with *kelch13* mutations if these have indeed originated outside Africa. We found that *kelch13* mutants in Africa do not possess similar flanking haplotypes to those found in SEA (Figure 2). This suggests that mutations observed in Africa do not have a common origin with those in Asia, and are likely to have emerged independently.

**Figure 2.**
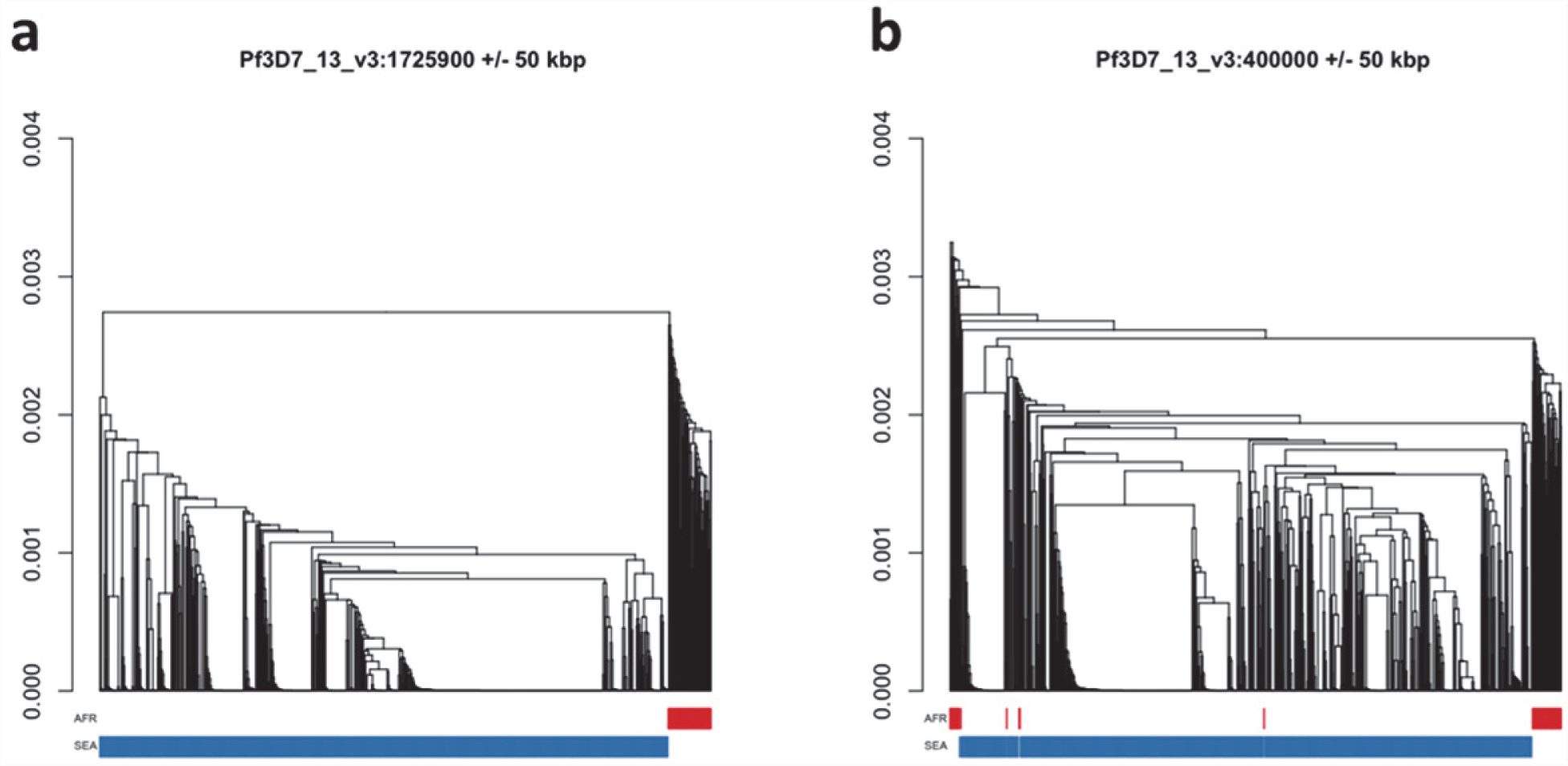
Different origin of African and Asian mutations. (a) Hierarchical clustering tree of *kelch13* mutants in SEA (n=741) and Africa (n=56), computed from pairwise genetic distances over a 100 kbp window centred at the *kelch13* locus. Haplotypes surrounding *kelch13* in SEA mutants (blue) form a separate tree from those in African mutants (red), suggesting that the latter have originated independently. (b) An analogous tree using a different 100 kbp region (centred at 400 kbp in the same chromosome) shows considerably less marked separation between mutants in the two continents.

### Across the genome there are many more rare variants in Africa

Historical demographic changes such as population expansions and bottlenecks (Tanabe et al. 2010), and epidemiological and environmental factors (Prugnolle et al. 2010) are highly influential forces that shaped the allele frequency spectra of *P. falciparum* populations across the globe (Nielsen et al. 2009; Manske et al. 2012). In order to properly contextualize the numbers and frequencies of *kelch13* mutations, it is therefore important to characterize genomic variation patterns in different geographical regions.

One of the most striking features of this dataset is the high number of rare variations in the high-quality SNP list. At more than half of all polymorphic sites, the minor allele was only carried by a single sample (referred to as *singletons*, n=330,785 or 36%) or by two samples (*doubletons*, n=214,179 or 23%), often in heterozygous calls. By contrast, only 13,288 polymorphisms (1.4%) had a minor allele in ≥5% of samples. Rare alleles, however, are not evenly distributed geographically. There is a large excess of polymorphisms with minor allele frequency (MAF) below 0.1% in Africa (72% of all SNPs, vs. 18% in SEA), while numbers in the two regions are similar for SNPs with MAF>1% (2% of all SNPs, Figure 3a). Rare variations in Africa are not confined to a limited set of highly variable genes, but evenly distributed across the genome, as attested by the distribution of variants across all genes: SNP density in Africa (median=67 SNPs/kbp, interquartile range=51-84) is approximately 3.9 times higher than in SEA (median=17, IQR=13-22, *P*< 10^-16^, Figure 3b). Very similar ratios are estimated in both non-synonymous (Africa median SNP density=43, IQR=27-58; SEA median=11, IQR=7-15) and synonymous variants (Africa median SNP density=25, IQR=20-30, SEA median=6, IQR=4-8). Accordingly, we found virtually identical distributions of the ratio of non-synonymous to synonymous mutations (N/S ratio) in the two regions (Figure 3c). This suggests that the huge disparity in SNP density between the two regions is more likely to be the result of different demographic histories and epidemiological characteristics, such as changes in effective population size (Joy et al. 2003), rather than the product of different selective constraints.

**Figure 3.**
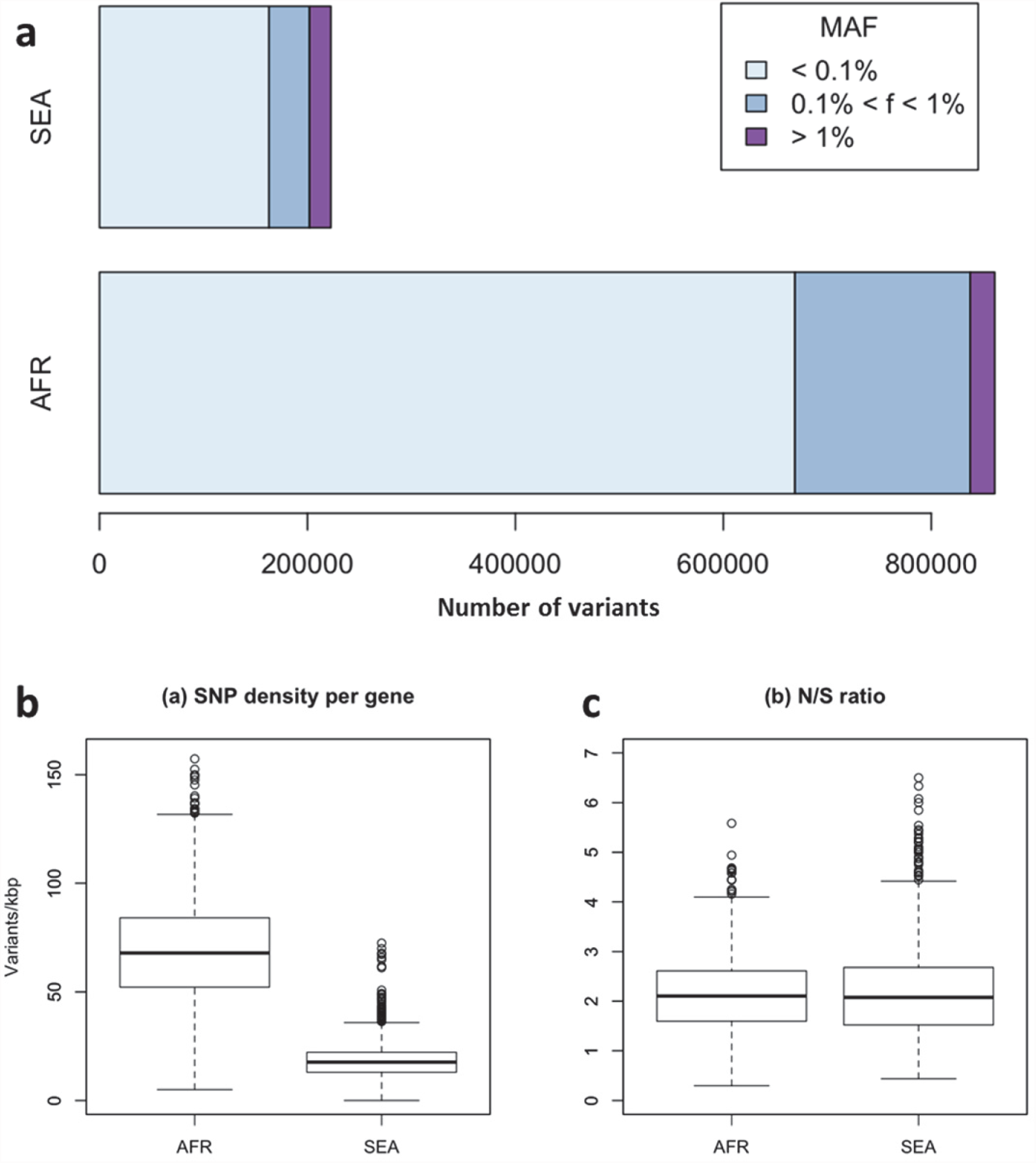
Number and density of variants in Africa and Southeast Asia. (a) Number of variants in Africa (AFR) and in SEA: while the number of high-frequency variations is consistent between the two regions, samples from Africa possess an excess of low-frequency variations and dominate the total number of variants discovered. (b) SNP density per gene: for each gene, the number of variants found in the two regions is normalized by the length of the coding region (in kbp). African samples have on average 3.9 times more mutations than parasites from SEA. (c) Non-synonymous / synonymous ratio per gene: ratios of non-synonymous to synonymous mutations found per gene are similar in the two regions. To reduce the effect of small numbers artefacts, in both analysis only genes with at least 10 SNP were considered.

In summary, we observe many more rare variants in Africa than in SEA; however we expect N/S ratios to be similar in these two regions in genes that are not subjected to selective pressures.

### Comparing *kelch13* with other parts of the genome

The density of *kelch13* synonymous variations in the two continents is roughly consistent with that observed in the rest of the genome (Africa: 28 SNPs/kbp, SEA: 6; Figure 4), which is expected since synonymous changes are less likely to be affected by selection. The excess of African non-synonymous mutations in the upstream region is also consistent with expectations (Africa: 44 SNPs/kbp, SEA: 16). In contrast, non-synonymous polymorphisms in the resistance domains show a reversal of this relationship: SEA parasites possess about 20% more mutations than African ones. In addition, all but two non-synonymous mutations in Africa are at very low frequency (mostly singletons and doubletons), while in SEA more than half of the changes are observed in >2 samples (Table 4).

**Figure 4.**
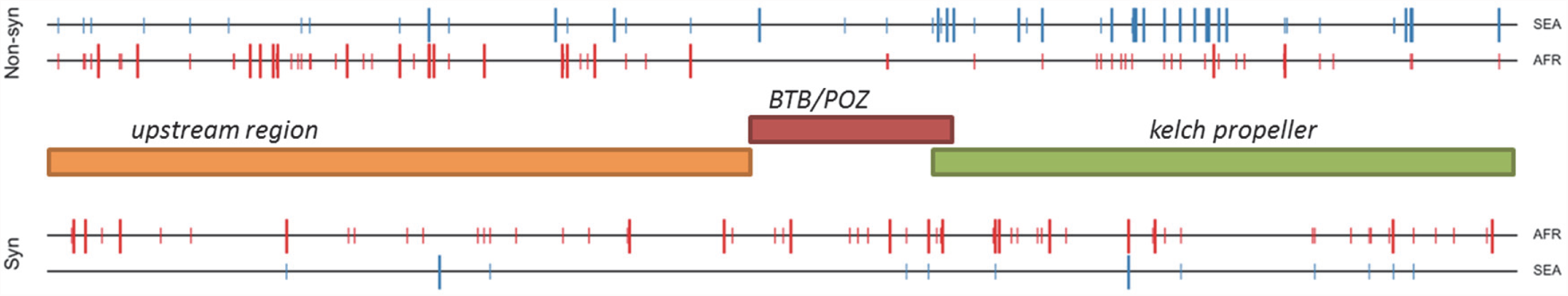
Structure of *kelch13* and positioning of mutations in Africa and Southeast Asia. The positions of *kelch13* polymorphisms observed in Africa (red) and SEA (blue) are shown. Coloured rectangles describe the extents of the resistance domains and upstream region, with the locations of non-synonymous changes indicated above, and that of synonymous changes below. Short lines represent singleton/doubleton mutations, while longer lines represent more common mutations.

**Table 4.** Frequency of the non-synonymous mutations in the *kelch13* resistance domains. Counts of non-synonymous mutations in the two conserved domains of *kelch13* are shown for each geographical region, stratified by the number of samples in which they are observed.

|  | AFR | SEA | SAS | OCE | SAM |
| --- | --- | --- | --- | --- | --- |
| *1-2 samples* | 24 | 13 | 1 | 1 | 0 |
| *3-5 samples* | 1 | 7 | 0 | 0 | 0 |
| *>5 samples* | 1 | 14 | 0 | 0 | 0 |

At a first approximation, these observations are consistent with a high number of non-synonymous changes that have risen in frequency in SEA parasites because of their association with artemisinin resistance, and a low number of mostly rare alleles in Africa, where artemisinin has been introduced more recently and resistance is yet to be reported.

### Comparing *kelch13* to other highly conserved genes

Although the function of *kelch13* is as yet unclear, an alignment of its homologous gene sequences in eight *Plasmodium* species shows that the resistance domains are part of a highly conserved region (Supplementary Figure 1), suggesting a crucial role in parasite fitness. A reconstruction of ancestral alleles from this alignment suggests that *P. falciparum* accumulated only five conservative amino acid changes in the *kelch13 propeller* domain since diverging from other species 55 Myr ago (Escalante and Ayala 1995) (Supplementary Table 1). Given this extreme level of conservation, non-synonymous polymorphisms may appear surprisingly numerous in the present dataset, both in SEA (n=34) and in Africa (n=26). Such elevated numbers may be produced by selection processes; alternatively, they may be present in a large neutrally-evolving population, in which low-frequency variations continually emerge, but are can only be detected for a brief span of time before they are removed by genetic drift and/or purifying selection. The question is, then, whether neutral evolution can account for the pattern of *kelch13* mutations observed here.

To answer this question, we compared patterns of *kelch13* mutations to those in the rest of the *P. falciparum* genome. Since fewer non-synonymous mutations are expected in more conserved genes, we applied *genomic calibration*, i.e. we stratified these analyses by evolutionary conservation. Each gene was assigned a *conservation score* determined from a sequence alignment of the *P. falciparum* gene with its *P. chabaudi* homologue, using a substitution matrix corrected for the AT bias in the *Pf* genome (Brick and Pizzi 2008). *P. chabaudi* was chosen as representative of rodent plasmodia, the group most differentiated from *P. falciparum*. A genome-wide non-linear negative correlation between gene conservation and N/S ratio is clearly observable; this trend is almost identical in the two populations (Figure 5a). Although *kelch13* did not diverge significantly from this relationship in Africa (P=0.2), its N/S ratio in SEA was the highest observed at its level of conservation, far exceeding the expected ratio (P<0.001). Accordingly, *kelch13* showed the most significant difference in N/S ratios between Africa and SEA genome-wide (3.7-fold, P=2x10^-4^ by Fisher’s exact test, Figure 5b). Such unusually high N/S ratio in SEA parasites is mainly due to an excess of high frequency non-synonymous variations (Supplementary Figure 2), suggesting that multiple independent origins of artemisinin resistance (Miotto et al. 2015; Takala-Harrison et al. 2015) have produced an unusually large number of common non-synonymous mutations.

**Figure 5.**
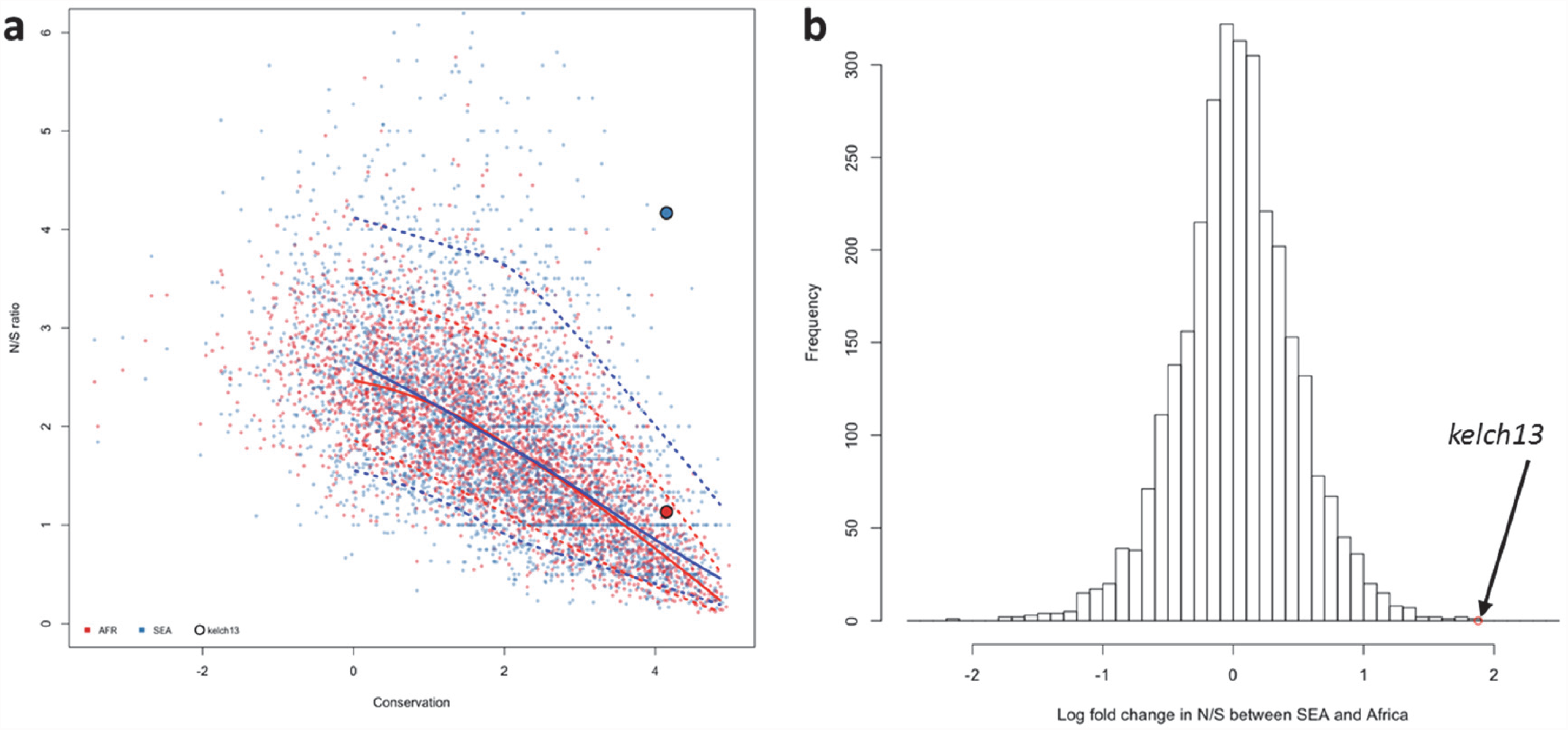
Genome-wide analysis of N/S ratio. (a) For each gene with more than 2 synonymous or non-synonymous SNPs, the N/S ratio in Africa (red points) and in SEA (blue points) are plotted against the conservation score (derived from a CCF53P62 matrix) of the gene coding sequence. The *kelch13* gene is represented by larger circles. For each region, a solid line show median values, while dotted lines delimit 95% of the genes at varying levels of conservation. (b) Histogram showing the distribution of the ratio of N/S ratios in SEA and Africa, for all genes with ≥2 either synonymous or non-synonymous SNPs on each region. An arrow shows the placement of *kelch13*.

From this analysis we conclude that the high prevalence of *kelch13* non-synonymous variants in SEA is not explainable by neutral evolution, but is consistent with selection of artemisinin-resistance alleles. In Africa, on the other hand, the observed non-synonymous changes appear to constitute a “physiological” level of variation consistent with a population rich in low-frequency alleles.

### In Southeast Asia there are more radical substitutions in *kelch13*

The different *kelch13* mutation repertoires in Africa and in SEA raise the question of whether these sets of mutations have different structural and functional properties. While there is high conservation across the whole of the resistance domains, it is unlikely that all possible amino acid changes have the same functional relevance or that they all carry the same fitness cost for parasites. Although direct measures of functional relevance are not yet available, and the exact function of *kelch13* is hitherto unknown, we can make statistical comparisons of some properties of the observed changes, in at least two respects. First, assuming that *kelch13* function is conserved across *Plasmodium* species, we can assess the strength of evolutionary constraints at any given position by examining whether amino acid substitutions between species are conservative or radical. Second, given that *kelch* proteins have been shown in other species to have an adapter role, with key binding sites defined by the arrangement of hydrophobic β-strands in the propeller domain, we can assess the hydrophobicity of the observed mutations, which should be informative of their functional importance.

We characterized all changes in the *kelch13* resistance domains with a conservation score derived from a substitution matrix specific to AT-rich genomes (Brick and Pizzi 2008), and assigned a hydrophobicity score to each site, estimated from the Kyte-Doolittle (KD) hydropathicity score (Kyte and Doolittle 1982). The five putative derived alleles that have emerged in the *P. falciparum kelch13 propeller* domain since its divergence from other *Plasmodia* are all conservative changes at hydrophilic sites (Figure 6a). The nature of mutations common in Africa is broadly consistent with this conservative history of change (Figure 6b and Supplementary Table 2). Polymorphisms in SEA parasites, on the other hand, show a contrasting pattern of changes that are more radical than those in Africa (P=10^-3^) and more commonly found at hydrophobic sites, which may support their role in modulating the substrate binding properties of the propeller (Figure 6c and Supplementary Table 2).

**Figure 6.**
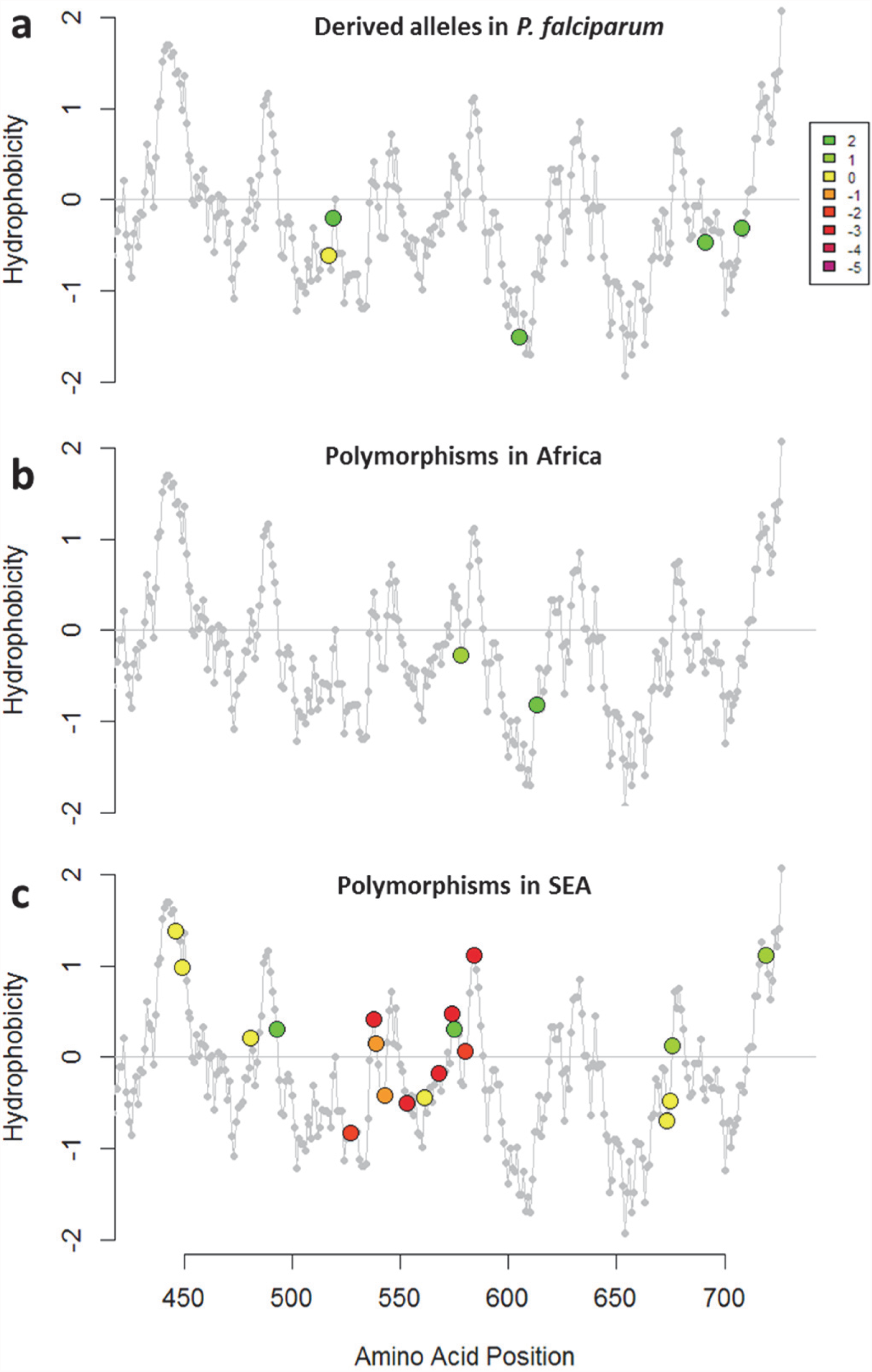
Characterization of kelch13 mutations in Africa and Southeast Asia. In these plots, all *kelch13* amino acids in the propeller domains, downstream of position 430, are plotted against their Kyte-Doolittle hydrophobicity score (KD, gray). Circle symbols in plot (a) derived amino acid alleles in *falciparum*, coloured according to the conservation score against the ancestral allele (Supplementary Table 1), derived from the CFF53P62 substitution matrix; lower values indicate more radical substitutions. The remaining plots use the same colouring scheme to show polymorphisms observed in ≥3 samples in Africa (b) or in SEA (c).

Detailed mapping against the secondary structure of the propeller domain suggests that the polymorphisms found in SEA parasites occur in different blades, preferentially at positions proximal to the first and second β-strand of the propeller’s blades (Figure 7a). These two β-strands provide local hydrophobicity peaks for each blade in a periodic pattern within the *kelch* propeller domain, presumably critical to the binding properties of the molecule (Figure 7b). Although the limited numbers do not support statistical inferences, we note that SEA polymorphisms abound in blades 3 and 4, and are almost absent in blade 5, where one of the three common African mutations is found. Taken together, the above results suggest that the resistance domains have long been under very strong evolutionary constraints, and that the number and nature of the changes observed in African parasites is consistent with these constraints, once we discard the abundant rare alleles expected in such a large population. In contrast, mutations in SEA parasites are not only far more numerous than expected, but they produce radical changes at sites that are likely to be important determinants of the binding properties of the *kelch13* protein. These findings bolster the hypothesis that these changes modulate *kelch13* functionality, and are likely agents of phenotypic change.

**Figure 7.**
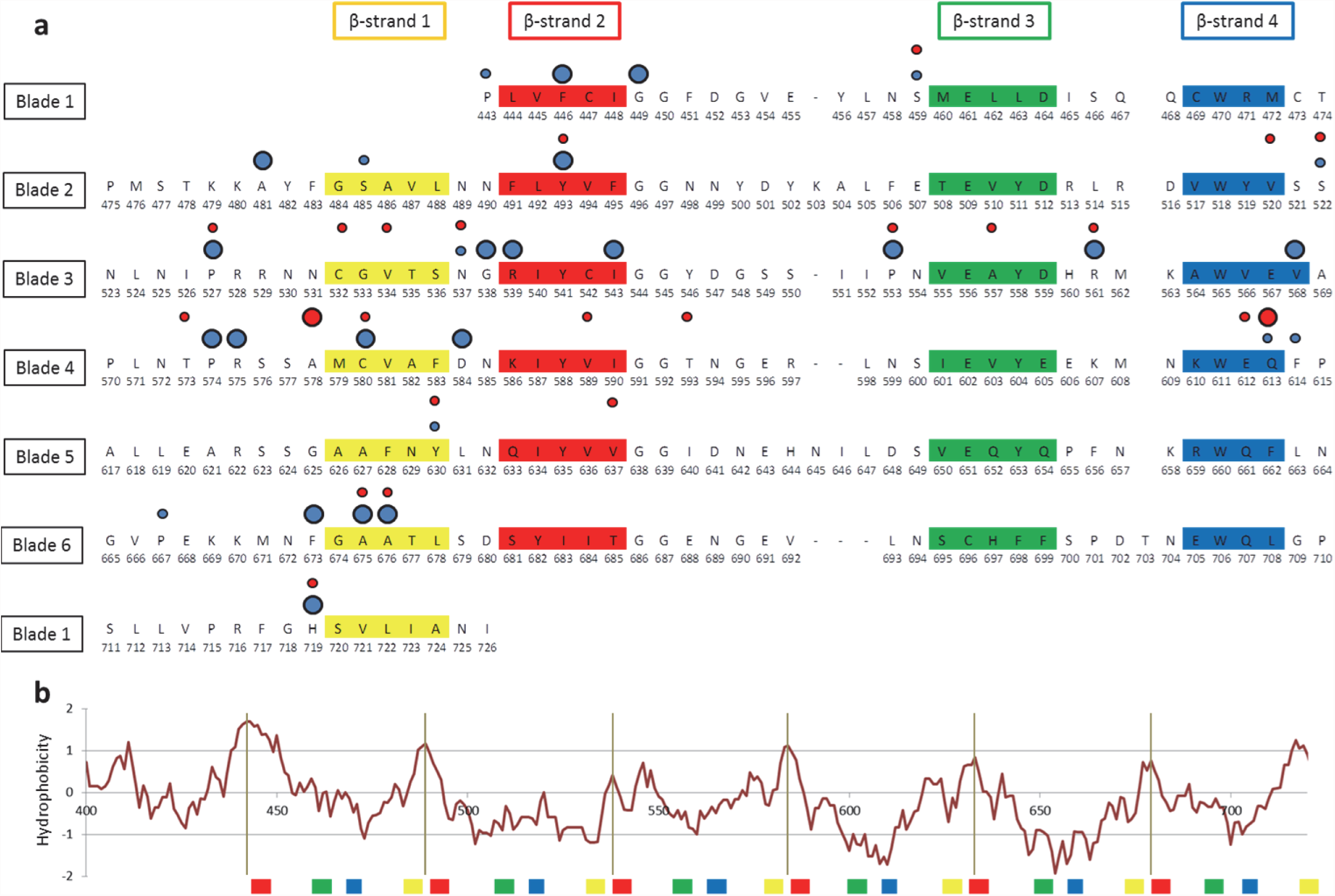
Structure of the kelch13 propeller domain, showing the position of mutations in Africa and Southeast Asia. (a) The sequence of the *kelch13* propeller domain (amino acids 443-726) is organized according to its 6-blade tertiary structure, with the four β-strands characterizing each blade highlighted in colour. Polymorphisms observed in Africa (red) and SEA (blue) are shown by circles above the mutated position. Small circles indicate very rare mutations (singletons and doubletons), while larger circles are used for more frequent mutations. (b) Plot of hydrophobicity at positions in this domain (estimated by Kyte-Doolittle score) showing how hydrophobicity peaks (gray vertical lines) occur periodically in correspondence to β-strands 1 and 2. The location of β-strands is shown below the plot, using the same colour scheme as above.

### Genetic background

A recent study has shown that resistance-causing K13PBD mutations are significantly more likely to arise in parasites with a particular genetic background (Miotto et al. 2015). This predisposing genetic background is marked by specific SNP alleles of the genes encoding ferredoxin (*fd*), apicoplast ribosomal protein S10 (*aprs10*), multidrug resistance protein 2 (*mdr2*) and chloroquine resistance transporter (*crt*). Here we extend this analysis, confirming that this particular combination of variants is extremely common in the parts of Southeast Asia where artemisinin resistance is known to be established, and is absent from Africa and other regions sampled here (Table 5).

**Table 5.** Frequency of the genetic background alleles across the world. Frequency of the four genetic background alleles identified in Miotto et al. 2015 for each geographical region. For each SNP, we show mutation name; chromosome number; nucleotide position; and frequencies of the mutant allele in the various populations.

| Mutation | Chr | Pos | AFR | SAS | SEA | PNG | SAM |
| --- | --- | --- | --- | --- | --- | --- | --- |
| **arps10-V127M** | 14 | 2481070 | 0.0% | 0.0% | 59.4% | 0.0% | 0.0% |
| **fd-D193Y** | 13 | 748395 | 0.1% | 2.2% | 62.8% | 23.9% | 0.0% |
| **mdr2-T484I** | 14 | 1956225 | 0.1% | 5.7% | 64.2% | 0.4% | 0.0% |
| **crt-N326S** | 7 | 405362 | 0.8% | 28.2% | 68.6% | 0.1% | 0.0% |

## Discussion

This study demonstrates the value of genomic epidemiology in characterising the current wave of artemisinin resistance, which is problematic for conventional molecular epidemiology since new resistance-causing mutations are continually emerging on different haplotypic backgrounds. A key problem is to define the geographical origin of KBPD mutations, and we show that this can be solved by using genomic epidemiological data to analyse ancestral relationships between samples, thereby demonstrating that the KPBD mutations observed in Africa are of local origin.

Another important question is whether KPBD mutations are under positive selection in Africa, which is difficult to determine by standard haplotype-based methods because so many independent mutations are involved. This question is further complicated by marked geographical variation in normal levels of genetic diversity, i.e. there are many more rare variants in Africa than Southeast Asia, most likely due to the larger population size and other demographic factors (Manske et al. 2012). Here we address this question by comparing *kelch13* against other genes in the same samples, a process that we refer to as genomic calibration. We show that for most genes the ratio of non-synonymous to synonymous mutations is relatively constant across geographical regions, despite geographic differences in genetic diversity, and that this ratio is correlated with the level of sequence conservation across different *Plasmodium* species. When calibrated against other genes with the same level of cross-species conservation, allowing for geographical differences in the overall level of genetic diversity, *kelch13* shows a marked excess of non-synonymous substitutions in Southeast Asia, but appears normal in Africa. Moreover, KPBD mutations causing radical amino acid changes at highly conserved positions are found at relatively high frequency in Southeast Asia but remain at very low frequency in Africa. Taken together, these findings indicate that non-synonymous KBPD mutations are undergoing strong evolutionary selection in Southeast Asia, whereas those seen in Africa have originated locally and most likely reflect normal standing variation.

These findings have practical implications for the prevention of artemisinin resistance in Africa, where there is evidently a deep reservoir of standing genetic variation that could potentially allow resistance to emerge rapidly as the levels of selective pressure increase. In most parts of Africa, the selective pressure of artemisinin is probably relatively low at present, for several reasons. Artemisinin has been widely used in Southeast Asia for over two decades, whereas its usage in Africa is more recent, and it is estimated that only 20% of infected African children currently have access to frontline treatment ACT medication (World Health Organization 2014). Another factor is that people living in regions of high malaria endemicity acquire partial immunity and asymptomatic infection, so that there is a large reservoir of parasites in Africa that are not exposed to antimalarial drugs because asymptomatic individuals do not seek treatment (Hastings 2003). The situation could change dramatically as malaria control efforts are intensified, and it will be vital to monitor the effects of major interventions on the emergence of resistance, particularly in African countries that have already achieved relatively low levels of malaria transmission. There also needs to be vigilance for significant changes in other parts of the genome, e.g. at the *fd, arps10*, *mdr2* and *crt* loci that appear to be predisposing factors for the emergence of resistance-causing KPBD mutations in Southeast Asia. Another concern is that growing resistance to ACT partner drugs, now emerging in Southeast Asia (Saunders et al. 2014). may spread to Africa and lead to increased selective pressure for artemisinin resistance there.

Previous waves of resistance to frontline antimalarial drugs, i.e. chloroquine and sulfadoxine-pyrimethamine, included localised emergences; but at a global level these were dominated by hard sweeps of specific haplotypes originating in Southeast Asia (Mita et al. 2009). Although we still know relatively little about the functional properties of different KPBD mutations, it is clear that some are more successful than others, e.g. the C580Y allele has emerged at multiple locations in Southeast Asia and Africa, and a specific C580Y haplotype is approaching fixation in large parts of Western Cambodia (Miotto et al. 2015). The high level of sequence conservation of KPBD across *Plasmodium* species indicates that mutations in these domains incur fitness costs, making mutant parasites less likely to survive. However, these fitness costs may be compensated by other genetic variants, either in *kelch13* or elsewhere in the genome, that are likely to accumulate in the course of evolution. The danger is that, as a result of this continuing evolutionary process, parasites in Southeast Asia will progressively acquire higher levels of artemisinin resistance (World Health Organization 2014) coupled with strong biological fitness and the ability to propagate across a wide range of vector species. Under these circumstances, the current soft sweep of artemisinin resistance could give way to a pervasive hard sweep with potentially disastrous consequences.

These findings demonstrate the utility of applying genomic epidemiology to identify features of parasite demography and evolution that affect how drug resistance spreads. Future strategies to combat resistance will require better understanding of the evolutionary consequences of malaria control interventions, e.g. how the selective advantage of a resistance allele is counterbalanced by its fitness cost under different control regimes and in different geographical settings. It is now possible to approach this problem prospectively, by conducting systematic spatiotemporal sampling and genome sequencing of the parasite population as an integral part of public health interventions to prevent resistance spreading.

## Acknowledgements

This study was conducted by the MalariaGEN *Plasmodium falciparum* Community Project, and was made possible by clinical parasite samples contributed by partner studies, whose investigators are represented in the author list. An extended list of acknowledgments, listing individuals who contributed to partner studies or to the MalariaGEN Resource Centre, is available in the Supplementary Materials. The sequencing, analysis, informatics and management of the Community Project are supported by the Wellcome Trust through Sanger Institute core funding (077012/Z/05/Z; 098051) and a Strategic Award (090770/Z/09/Z) and the Medical Research Council through the MRC Centre for Genomics and Global Health (G0600230). This work was supported in part by the Intramural Research Program of the NIAID, NIH and by the Bill and Melinda Gates Foundation.

## Author Contributions

AA, CA, LA, TJCA, VA, TA, EA, SA, GAA, VB, AB, MFB, SB, TB, OB, PCB, KC, DJC, AC, NPD, AD, CDo, AMD, CDr, PD, DFE, TGE, RMF, MAF, CIF, TTH, AVOH, MI, DI, PL, CL, JMa, KM, MM, VM, OAM, JMo, IM, PKM, PNN, FN, RN, AN, HO, AO, MO, JO, APP, CVP, RNP, SP, MR, PR, LR, DS, AS, PS, ST, TNT, VT, FV, NJW, HY carried out clinical studies and/or laboratory work to obtain *P. falciparum* samples for sequencing. SC, DMe, ED, MS developed and implemented methods for sample processing and sequencing library preparation. RA, OM, JA, HMM, RP developed software tools and methods. CM, DJ, HMM, JS managed data production pipelines. RA, OM, VJC, RG, DMu, JCR, GM, KR, AM, BM, DPK contributed to study design and management. RA, OM, CW, JA, IS, DPK performed data analyses. RA, OM, DPK drafted the manuscript which was reviewed by all authors.

## Author Information

All sequence data are available online at the European Nucleotide Archive (ENA). The authors declare no competing financial interests.

## Methods

### Ethical Approval

All samples in this study were derived from blood samples obtained from patients with P. falciparum malaria, collected with informed consent from the patient or a parent or guardian. At each location, sample collection was approved by the appropriate local and institutional ethics committees. A full list of these committees is given in the Supplementary Materials.

### Sample Preparation, Sequencing and Genotyping

DNA was extracted directly from blood samples taken from patients at admission time, after leukocyte depletion to minimize human DNA. Leukocyte depletion was achieved by CF11 filtration in most samples (Venkatesan et al. 2012), or alternatively by Lymphoprep density gradient centrifugation (Axis-Shield) followed by Plasmodipur filtration (Euro-Diagnostica)(Auburn et al. 2011) or by Plasmodipur filtration alone. Genomic DNA was extracted using the QIAamp DNA Blood Midi or Maxi Kit (Qiagen), and quantities of human and Plasmodium DNA were determined by fluorescence analysis using a Qubit instrument (Invitrogen) and multi-species quantitative PCR (Q-PCR) using the Roche Lightcycler 480 II system, as described previously (Manske et al. 2012). Samples with >50 ng DNA and <80% human DNA contamination were selected for sequencing on the Illumina HiSeq platform following the manufacturer’s standard protocols (Bentley et al. 2008). Paired-end sequencing reads of length 200-300bp were obtained, generating approximately 1Gbp of read data per sample. All short read sequence data have been deposited in the European Nucleotide Archive (http://www.ebi.ac.uk/ena/data/search/?query=plasmodium), and metadata will be released at the time of publication.

Polymorphism discovery, quality control and sample genotyping followed a process described elsewhere (Manske et al. 2012). Short sequence reads from 3,411 *P. falciparum* samples included in the *MalariaGEN Plasmodium falciparum Community Project* (https://www.malariagen.net/node/44) were aligned against the *P. falciparum* 3D7 reference sequence V3 (ftp://ftp.sanger.ac.uk/pub/pathogens/Plasmodium/falciparum/3D7/3D7.latest_version/version3/), using the bwa program(Li and Durbin 2009) (http://bio-bwa.sourceforge.net/) as previously described(Manske et al. 2012), to identify an initial global set of 4,305,957 potential SNPs. This list was then used to guide stringent re-alignment using the SNP-o-matic algorithm (Manske and Kwiatkowski 2009), to reduce misalignment errors. The stringent alignments were then examined by a series of quality filters, with the aim of removing alignment artefacts and their sources. In particular, the following were removed: a) non-coding SNPs; b) SNPs where polymorphisms have extremely low support (<10 reads in one sample); c) SNPs with more than two alleles, with the exception of loci known to be important for drug resistance, which were manually verified for artifacts; d) SNPs where coverage across samples is lower than the 25^th^ percentile and higher than the 95^th^ percentile of coverage in coding SNPs (these thresholds were determined from artifact analysis); e) SNPs located in regions of relatively low uniqueness (Manske et al. 2012); f) SNPs where heterozygosity levels were found to be inconsistent with the heterozygosity distribution at the SNP’s allele frequency; and g) SNPs where genotype could not be established in at least 70% of the samples. These analyses produced a final list of 926,988 high-quality SNPs in the 14 chromosomes of the nuclear genome, whose genotypes were used for analysis in this study. All samples were genotyped at each high-quality SNP by a single allele, based on the number of reads observed for the two alleles at that position in the sample. At positions with fewer than 5 reads, the genotype was undetermined (no call was made). At all other positions, the sample was determined to be *heterozygous* if both alleles were each observed in more than 2 reads; otherwise, the sample was called as homozygous for the allele observed in the majority of reads. For the purposes of estimating allele frequencies and genetic distances, a within-sample allele frequency (*f*_*w*_) was also assigned to each valid call. For heterozygous calls, *f*_*w*_ was estimated as ratio of non-reference read count to reference read count; homozygous calls were assigned *f*_*w*_=0 when called with the reference allele, and *f*_*w*_=1 when called with the non-reference allele.

### Frequency estimation and clustering

For a given population *P*, we estimated the *non-reference allele frequency* (*NRAF*) at a given SNP as the mean of the within-sample allele frequency (*f*_*w*_) for all samples in *P* which have a valid genotype at that SNP. The *minor allele frequency* (*MAF*) at is the computed as min(*NRAF*, (1 – *NRAF*)). To investigate the origin of the *kelch13* mutations in Africa, we clustered by genetic distance all the samples carrying a non-synonymous mutation in the resistance domains. The clustering is expected to group together samples with similar haplotypes and hence likely to have the same origin. The topology of the resulting tree was then compared to the one obtained using the same method to a region of the genome not associated to artemisinin-resistance. To this aim, we considered two different regions of the genome. One centred on *kelch13* and extending 50kbp both sides (Pf3D7_13_v3:1,775,900-1,675,900), and one on the same chromosome but centred upstream (Pf3D7_13_v3:350,000-450,000) serving as control. This control region lies outside of the main haplotypes associated to *kelch13* (Miotto et al. 2015). Different window sizes led to similar results. The genotype of *kelch13* was derived from read counts at non-synonymous SNP in the *kelch13* resistance domains using the same procedure described previously (Miotto et al. 2015). We computed an *N*x*N* pairwise distance matrix, where *N* is the number of samples. Each cell of the matrix contained an estimate of genetic distance between the relevant pair of samples, obtained by summing the pairwise distance, estimated from within-sample allele frequency (*f*_*w*_), at each SNP in the 100kbp window considered. When comparing a pair of samples *s*_A_ and *s*_B_ at a single SNP *i* where a genotype could be called in each sample, with within-sample allele frequencies *f*_*A*_ and *f*_*B*_ respectively, the distance *d*_*AB*_ was estimated as *d*_*AB*_ = *f*_*A*_ (1- *f*_*B*_) + *f*_*B*_ (1- *f*_*A*_). The genome-wide distance D_AB_ between the two samples is then calculated as

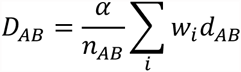

where *n*_AB_ is the number of SNPs where both samples could be genotyped, *w*_*i*_ is an LD weighting factor (see below) and a is α scaling constant, equal to 70% of the number of coding positions in the genome (since our genotyping covers approximately 70% of the coding genome). The exact value of α is uninfluential towards the analyses conducted in this study. The LD weighting factor, which corrects for the cumulative contribution of physically linked polymorphisms, was computed at each SNP *i* with *MAF* ≥ 0.1 in our sample set, by considering a window of *m* SNPs (*j* = 0.. *m*) centred at *i*. For each *j*, we computed the squared correlation coefficient *r2ij* between SNPs *i* and *j*. Ignoring positions *j* where where r2*ij* < 0.1, the weighting *w*_*i*_ was computed by

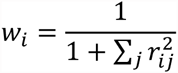

A tree was then built using a standard hierarchical clustering method (`hclust`) implemented in the R `stats` package (http://stat.ethz.ch/R-manual/R-patched/library/stats/html/00Index.html) using the Ward’s minimum variance criterion. A similar pairwise distance matrix, but using the whole genome, was also used to produce a neighbour-joining tree using the nj implementation in the R ape package.

### Conservation scoring and ancestral allele analysis

We analyzed homologous protein sequences of *kelch13* genes for seven *Plasmodium* species for which high-quality sequence data were available: *P. falciparum*, *P. reichenowi*, *P. vivax*, *P. knowlesi*, *P. yoelii*, *P. berghei*, and *P. chabaudi*. The sequences were retrieved from the OrthoMCL cluster ORTHOMCL894 in GeneDB (http://www.genedb.org/), and a multiple alignment was obtained using ClustalW (Larkin et al. 2007) at default settings. In turn, we considered each pair alignment of *P. falciparum* with one of the remaining species, assigning a *substitution score* to each *kelch13* amino acid position, derived from the CCF53P62 substitution matrix. Although this matrix was chosen due to its suitability for AT-rich codon biases (Brick and Pizzi 2008), we found that use of the more commonly used BLOSUM62 matrix (Henikoff and Henikoff 1992) did not have a significant effect on the results. Finally, each amino acid position was assigned a *conservation score* for the pair alignment, equal to the mean of the substitution scores in a 9-residue window centered at that position.

To reconstruct putative *ancestral* and *derived* alleles in the propeller domain, we catalogued all polymorphic positions in the multiple sequence alignment. We organized the seven species into three groups by similarity: Laverania (*P. falciparum*, *P. reichenowi*), primate Plasmodia (*P. vivax*, *P. knowlesi*) and rodent Plasmodia (*P. yoelii*, *P. berghei*, and *P. chabaudi*), and observed that at each position, only one of the groups presented an allele different from that in the remaining groups. This group-specific allele was labelled as a putative *derived* allele, and the alternative allele as *ancestral*. (see Supplementary Table 1). Rodent species were found to carry the highest number of derived alleles, and therefore deemed to be good comparators for genome-wide conservation scoring. *P. chabaudi* was selected as a representative species in this group, and used for subsequent comparative analyses.

We estimated a *gene conservation score* for every *P. falciparum* gene for which a *P. chabaudi* orthologue sequence could be obtained from PlasmoDB (http://www.plasmodb.org/). The details of the method are described elsewhere (Gardner et al. 2011). Briefly, alignments of orthologous protein sequences were performed using ClustalW (Larkin et al. 2007) at default settings, and each amino acid position was assigned a CCF53P62 substitution score (see above). The gene conservation score assigned was equal to the mean substitution score for all amino acid positions in the gene.

### Hydrophobicity analysis

Each amino acid position in the *kelch13* was assigned a *hydrophobicity score*, estimated by computing the mean of the Kyte-Doolittle index in a 14-residue window centered at the position, using the protein sequence translated from the 3D7 reference sequence.

## Supplementary Materials

### Full Acknowledgements

The authors would like to thank the following individuals, who contributed to partner studies or to the MalariaGEN Resource Centre, making this study possible.

- **Ghana**: James Abugri, Nicholas Amoako
- **Kenya**: Steven M. Kiara, John Okombo
- **Madagascar**: Rogelin Raherinjafy, Seheno Razanatsiorimalala
- **U.S.A.**: Hongying Jiang, Xin-zhuan Su

### Supplementary Figures

**Supplementary Figure 1.**
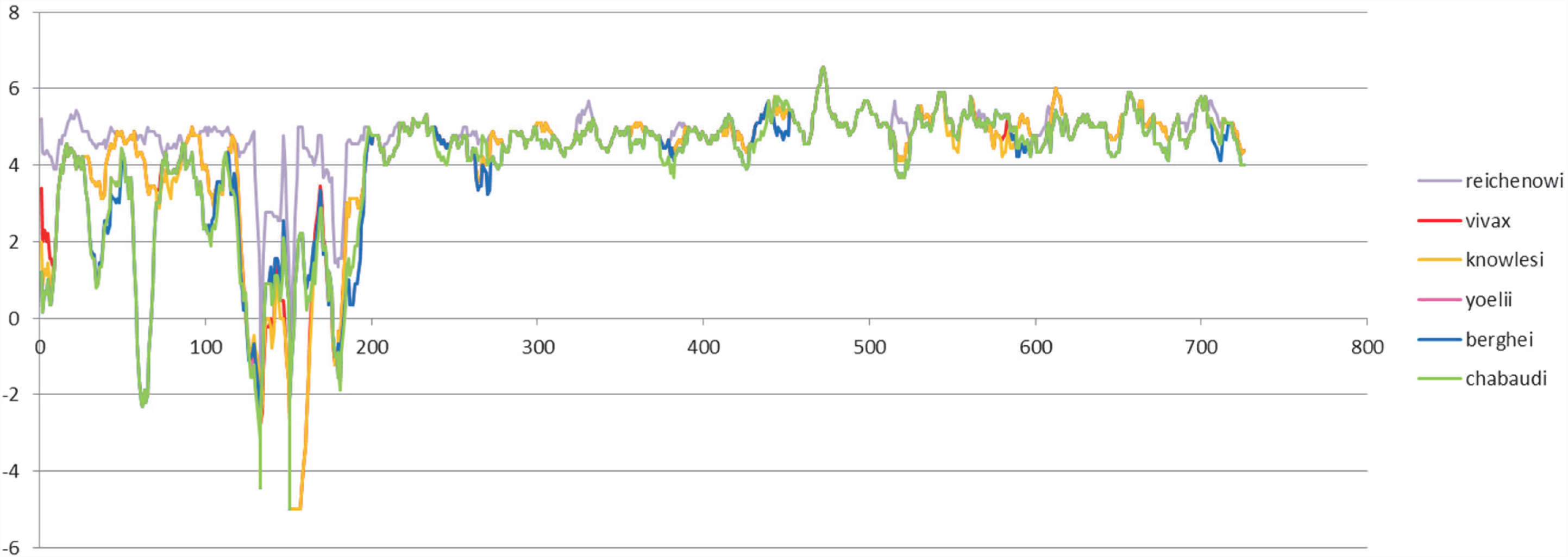
Conservation of *kelch13* sequence in *Plasmodium* spp. Conservation score across the 726 amino acid residues of *kelch13*, derived by applying a CCF53P62 matrix on alignments of the *P. falciparum* gene coding sequence with its homologues in six other *Plasmodium* species for which high-quality sequence data were available: *P. reichenowi*, *P. vivax*, *P. knowlesi*, *P. yoelii*, *P. berghei*, and *P. chabaudi* (see Methods). The region containing the resistance domains is highly conserved between species, whereas the upstream region is generally less conserved. Although the region below position 200 is less conserved, particularly in rodent species (*P. yoelii*, *berghei*, and *chabaudi*), there is remarkably high conservation across all species over the rest of the gene, which includes the two resistance domains (*BTB/POZ* and *propeller*).

**Supplementary Figure 2.**
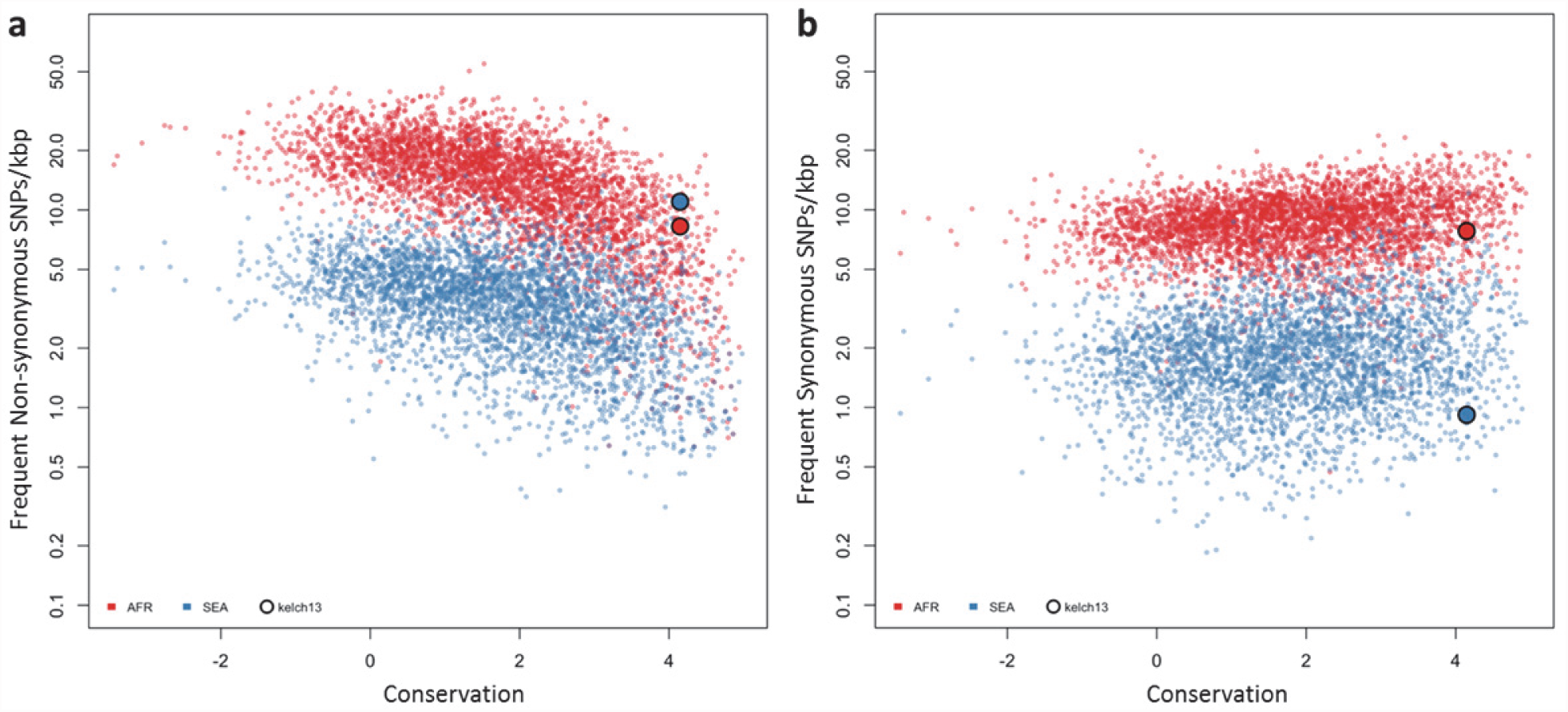
Genome-wide analysis of high-frequency SNP density. (a) Plot of the density of frequent (present in 3 or more samples) non-synonymous mutations in Africa (red) and SEA (blue) against conservation, for all genes with more than 2 synonymous or non-synonymous SNPs, showing that *kelch13* (square symbol) is consistent with other similarly conserved genes in Africa, but has excess non-synonymous polymorphisms in SEA. (b) An analogous plot for synonymous mutations also shows that *kelch13* follows the normal trend among genes in Africa, but has far fewer SNPs than expected in SEA.

### Supplementary Tables

**Supplementary Table 1.** *Kelch13 propeller* domain mutations in different *Plasmodium* species. Here we report amino acid allele differences in a multiple sequence alignments of *kelch13* homologues for seven species of *Plasmodium* parasites for which high-quality sequence data were available: *P. falciparum* (Pf), *P. reichenowi* (Pr), *P. vivax* (Pv), *P. knowlesi* (Pk), *P. yoelii* (Py), *P. berghei* (Pb), and *P. chabaudi* (Pc). The species formed three groups by similarity: Laverania (Pf, Pr), primate Plasmodia (Pv, Pk) and rodent Plasmodia (Py, Pb and Pc). An allele shared by all members of two different groups was identified as a putative ancestral allele. The table shows, for each position where at least one species exhibits a difference from the others: the amino acid position in the Pf *kelch13* sequence; the putative ancestral amino acid allele; the alleles in the various species (columns with heading listing multiple species show mutations common to those species); and a substitution score of the mutation, based on a CCF53P62 substitution matrix (see Methods). All substitution scores are ≥0, denoting conservative substitutions.

| Pf Position | Ancestral Allele | Pf,Pr | Pv,Pk | Pk | Py,Pb, Pc | Py,Pb | Pb | Pc | Substitution Score |
| --- | --- | --- | --- | --- | --- | --- | --- | --- | --- |
| 434 | F | . | . | . | . | . | . | Y | 2 |
| 447 | C | . | . | . | . | S | . | . | 1 |
| 448 | I | . | M | . | . | . | . | . | 2 |
| 517 | T | V | . | . | . | . | . | . | 0 |
| 519 | F | Y | . | . | . | . | . | . | 2 |
| 520 | V | . | . | . | L | . | . | . | 0 |
| 534 | V | . | . | . | I | . | . | . | 2 |
| 550 | S | . | . | C | . | . | . | . | 1 |
| 566 | V | . | . | . | I | . | . | . | 2 |
| 568 | V | . | I | . | . | . | . | . | 2 |
| 578 | A | . | S | . | . | . | . | . | 0 |
| 584 | D | . | . | E | . | . | . | . | 1 |
| 590 | I | . | . | . | . | V | . | . | 2 |
| 593 | T | . | . | . | A | . | . | . | 0 |
| 605 | D | E | . | . | . | . | . | . | 1 |
| 613 | Q | . | . | . | . | N | . | K | 0 |
| 648 | D | . | . | . | E | . | . | . | 1 |
| 666 | V | . | . | . | I | . | . | . | 2 |
| 676 | A | . | . | . | T | . | . | . | 0 |
| 691 | D | E | . | . | . | . | . | . | 1 |
| 708 | I | L | . | . | . | . | . | . | 0 |
| 711 | S | . | . | . | . | . | P | . | 0 |
| 723 | I | . | . | . | V | . | . | . | 2 |

**Supplementary Table 2.** Conservtion score of *kelch13* mutations. The table shows, for each non-synonymous *kelch13* mutation observed in the dataset, the number of samples carrying the mutation in Africa (AFR), in Southeast Asia (SEA), and a substitution score of the mutation, based on a CCF53P62 substitution matrix (see Methods); lower values indicate more radical substitutions. Mutations observed in Africa tend to have higher conservation score, whereas in SEA mutations tend to be more radical.

| Mutation | AFR | SEA | CCF53P62 |
| --- | --- | --- | --- |
| Q613E | 5 | 1 | 2 |
| Y630F | 2 | 1 | 2 |
| V637I | 2 |  | 2 |
| V589I | 2 |  | 2 |
| T573S | 2 |  | 2 |
| Y493H | 1 | 76 | 2 |
| I416V | 1 | 1 | 2 |
| T593S | 1 |  | 2 |
| V520I | 1 |  | 2 |
| I416M | 1 |  | 2 |
| F395Y |  | 1 | 2 |
| S522C | 2 | 1 | 1 |
| E612D | 1 |  | 1 |
| C532S | 1 |  | 1 |
| R575K |  | 3 | 1 |
| A578S | 18 |  | 0 |
| A676S | 2 | 3 | 0 |
| V534L | 2 |  | 0 |
| R561H | 1 | 24 | 0 |
| A675V | 1 | 18 | 0 |
| H719N | 1 | 8 | 0 |
| P527H | 1 | 5 | 0 |
| A557S | 1 |  | 0 |
| R539T |  | 63 | 0 |
| I543T |  | 34 | 0 |
| G449A |  | 7 | 0 |
| F446I |  | 7 | 0 |
| A481V |  | 4 | 0 |
| F673I |  | 3 | 0 |
| P667A |  | 2 | 0 |
| F614L |  | 1 | 0 |
| S485N |  | 1 | 0 |
| P443S |  | 1 | 0 |
| C580Y | 2 | 423 | -2 |
| D353Y |  | 4 | -2 |
| K438N |  | 1 | -2 |
| P553L | 2 | 24 | -3 |
| P441L |  | 27 | -3 |
| G538V |  | 19 | -3 |
| P574L |  | 12 | -3 |
| V568G |  | 6 | -3 |
| P667L |  | 2 | -3 |
| S459L | 2 | 2 | -4 |
| N537I | 1 | 1 | -4 |
| Q613L | 1 |  | -4 |
| D584V |  | 3 | -5 |

### Supplementary Text

#### Full List of Ethical Approval Authorities

This is the complete list of local and institutional committees that gave ethical approval for the partner studies: Comité d’Éthique, Ministère de la Santé, Bobo-Dioulasso, Burkina Faso; Navrongo Health Research Centre Institutional Review Board, Navrongo, Ghana; Kintampo Health Research Centre Institutional Ethics Committee, Kintampo, Ghana; Noguchi Memorial Institute for Medical Research Institutional Review Committee, University of Ghana, Legon, Ghana; Ghana Health Service Ethical Review Committee, Accra, Ghana; Gambia Government/MRC Joint Ethics Committee, Banjul, The Gambia; Comité d’Ethique National Pour la Recherche en Santé, Guinea; Ethics Committee of Faculté de Médecine, de Pharmacie et d’Odonto-Stomatologie, University of Bamako, Bamako, Mali; Ethical Review Committee, University of Ilorin Teaching Hospital, Ilorin, Nigeria; Institutional Review Board, Faculty of Health Sciences, University of Buea, Cameroon; Comité d’Ethique, Ecole de Santé Publique, Université de Kinshasa, Ministère de l’Enseignement Superieur, Universitaire et Recherche Scientifique, D. R. Congo; Comité National d’Ethique auprès du Ministère de la Santé Publique, Madagascar; Institutional Review Committee, Med Biotech Laboratories, Kampala, Uganda and Uganda National Council for Sciences and Technology (UNCST); College of Medicine Research Ethics Committee, University of Malawi, Blantyre, Malawi; KEMRI National Ethical Review Committee, Kenya; Medical Research Coordinating Committee of the National Institute for Medical Research, Tanzania; Ethical Review Committee, Bangladesh Medical Research Council, Bangladesh; Ethics Committee of the International Centre for Diarrheal Disease Research, Bangladesh; Institutional Ethical Review Committee, Department of Medical Research (Lower Myanmar); Ministry of Health, Government of The Republic of the Union of Myanmar; National Ethics Committee for Health Research, Ministry of Health, Phnom Penh, Cambodia; Ministry of Health National Ethics Committee For Health Research, Laos; Ethics Committee, Faculty of Tropical Medicine, Mahidol University, Bangkok, Thailand; Ethical Committee, Hospital for Tropical Diseases, Ho Chi Minh City, Vietnam; Eijkman Institute Research Ethics Commission, Jakarta, Indonesia; Institutional Review Board, Papua New Guinea Institute of Medical Research, Goroka, Papua New Guinea; Institutional Review Board, International Center for Medical Research and Training, Cali, Colombia; Institutional Review Board, Universidad Nacional de la Amazonia Peruana, Iquitos, Peru; Human Research Ethics Committee of NT Department of Health and Families and Menzies School of Health Research, Darwin, Australia; Institutional Review Board, New York University Medical School, NY, USA; Institutional Review Board, National Institute of Allergy and Infectious Diseases, Bethesda, MD, USA; Institutional Review Board, Walter Reed Army Institute of Research, Washington DC, USA; Ethics Review Committee, World Health Organization, Geneva, Switzerland; Ethics Committee of the Faculty of Medicine, Heidelberg, Germany; Ethics Committee of the Medical University of Vienna; Ethics Committee, London School of Hygiene and Tropical Medicine, London, UK; Oxford Tropical Research Ethics Committee, Oxford, UK.

